# The C-terminus of CFAP410 forms a tetrameric helical bundle that is essential for its localization to the basal body

**DOI:** 10.1101/2023.11.30.569397

**Authors:** Alexander Stadler, Laryssa De Liz, Heloisa B. Gabriel, Santiago Alonso-Gil, Robbie Crickley, Katharina Korbula, Bojan Žagrović, Sue Vaughan, Jack D. Sunter, Gang Dong

**Affiliations:** Max Perutz Labs, Vienna Biocenter, Medical University of Vienna, 1030 Vienna, Austria; Department of Biological and Medical Sciences, Oxford Brookes University, Oxford, OX3 0BP, UK; Departamento de Microbiologia, Imunologia e Parasitologia, Universidade Federal de Santa Catarina, Florianópolis, SC, Brazil; Department of Structural and Computational Biology, Max Perutz Labs, University of Vienna, Campus Vienna Biocenter 5, 1030, Vienna, Austria

**Keywords:** CFAP410, cilium, ciliopathies, flagellum, protein, structure

## Abstract

Cilia are antenna-like organelles protruding from the surface of many cell types in the human body. Defects in ciliary structure or function often lead to diseases that are collectively called ciliopathies. Cilia and flagella associated protein 410 (CFAP410) localizes at the basal body of cilia/flagella and plays essential roles in ciliogenesis, neuronal development, and DNA damage repair. It remains unknown how its specific basal body location is achieved. Multiple single amino acid mutations in CFAP410 have been identified in patients with various ciliopathies. One of the mutations, L224P, is located in the C-terminal domain (CTD) of human CFAP410 and causes severe <u>s</u>pondylo<u>m</u>etaphyseal <u>d</u>ysplasia, <u>ax</u>ial (SMDAX). However, the molecular mechanism for how the mutation causes the disorder remains unclear. Here, we report our structural studies on the CTD of CFAP410 from three distantly related organisms, *Homo sapiens, Trypanosoma brucei*, and *Chlamydomonas reinhardtii.* The crystal structures reveal that the three proteins all adopt the same conformation as a tetrameric helical bundle. Our work further demonstrates that the tetrameric assembly of the CTD is essential for the correct localization of CFAP410 in *T. brucei*, as the L224P mutation that disassembles the tetramer disrupts its basal body localization. Taken together, our studies reveal that the basal body localization of CFAP410 is controlled by the CTD and provide a mechanistic explanation for how the mutation L224P in CFAP410 causes ciliopathies in humans.

## Introduction

Cilia and flagella associated protein 410 (CFAP410) is coded by a gene that was mapped to chromosome 21 open reading frame 2 and thus named C21orf2 originally [1]. The gene was first thought as a candidate for a few genetic disorders based on its chromosomal location and the predicted mitochondrial localization of its coded protein [1]. Later mutational analysis excluded CFAP410 as the causative factor for those disorders and instead suggested it is a compelling candidate for retinal dystrophy [2, 3]. Immunofluorescence analysis showed that CFAP410 localizes to the basal body in mIMCD3 and hTERT-RPE1 cells as well as to the base of the connecting cilium in mouse photoreceptors, whereas genome-wide siRNA screen and biochemical studies suggest that it works together with NEK1 and SPATA7 as a functional module in ciliogenesis and DNA damage repair [4-6].

CFAP410 is also one of the genes implicated in amyotrophic lateral sclerosis (ALS), a fatal progressive neurodegenerative disease caused by the loss of motor neurons [7]. Moreover, the expression level of CFAP410 is substantially reduced in the brain of Down syndrome patients, implicating its function in neuronal development [8]. All these diversified functions of CFAP410 can be linked to its localization and function in cilia, as many neurological disorders are caused by defects in primary cilia [9].

CFAP410 exhibits significant sequence conservation across phyla and is present in all sequenced genomes of ciliates, including algae, protists, and animals. *Homo sapiens* CFAP410 (HsCFAP410) consists of 255 amino acids. Several single residue mutations of HsCFAP410 have been identified in patients with skeletal and/or retinal disorders including spondylometaphyseal dysplasia, axial (SMDAX) and retinal dystrophy with or without macular staphyloma (RDMS). Both disorders are genetically autosomal recessive and manifested as dysplasia of the ribs, vertebral bodies, ilia, proximal femora in skeletons, and retinitis pigmentosa in eyes [10, 11]. These clinically identified mutations include I35F, C61Y, R73P, Y107C, Y107H, V111M and L224P [4, 12-15]. However, it remains mysterious how these single-residue mutations of HsCFAP410 cause the deleterious defects due to a lack of high-resolution structures of the protein.

We found that CFAP410 consists of two folded modules, the N-terminal domain (NTD) and the C-terminal domain (CTD), which are connected via a long unstructured linker. Here we report our structural and biochemical characterizations on the CTD of three homologs of CFAP410 from *Homo sapiens, Trypanosoma brucei* and *Chlamydomonas reinhardtii.* We have determined high-resolution crystal structures for all three CTDs, which demonstrate that they all fold into a tetrameric helical bundle. We further revealed that the CTD is necessary for the targeting of CFAP410 to the basal body. The disease-causing L224P mutation both disrupts the tetrameric assembly and abolishes the basal body localization of CFAP410. Our work altogether explains how CFAP410 localizes to the basal body and why the single-residue mutation L224P causes ciliopathies.

## Results

### Crystal structures of the C-terminal domain of three CFAP410 orthologs

To investigate the three-dimensional structure of CFAP10, we chose homologs from three diverse species *H. sapiens*, *T. brucei* and *C. reinhardtii*. Our folding analyses suggested that all these CFAP410 orthologs contain two folded regions, including a globular domain at the N-terminus and a highly conserved small domain at the C-terminus (**Figure S1A-D**). The two domains are connected by a long disordered loop. Sequence alignments of homologs from various *Trypanosoma* species showed that both domains are highly conserved, whereas the connecting loop has very low homology and is variable in length (**Figure S1E**). Similar conservation patterns are also seen in homologs from mammals and algae (**Figure S2A-C**), and such pattern is also preserved across species (**Figure S2D**).

We attempted to crystallize the CTDs of *H. sapiens*, *T. brucei* and *C. reinhardtii* CFAP410 proteins. Multiple truncations were chosen based on sequence conservation and secondary structural predictions. Synthetic polypeptides of these truncations were further purified and subsequently used for crystallization trials. The CTD truncations that finally generated well-diffracting crystals were fragments covering residues 256-295, 212-246 and 213-246 of *H. sapiens*, *T. brucei* and *C. reinhardtii* CFAP410 proteins, respectively. The crystal structures were determined at 1.29-1.50 Å resolution (**Table 1**). The 2*F_o_-F_c_* electron density maps for all three structures show very high quality, with the side chains of almost all residues in these CTDs having been confidently built and refined (**Figure 1**).

**Figure 1.**
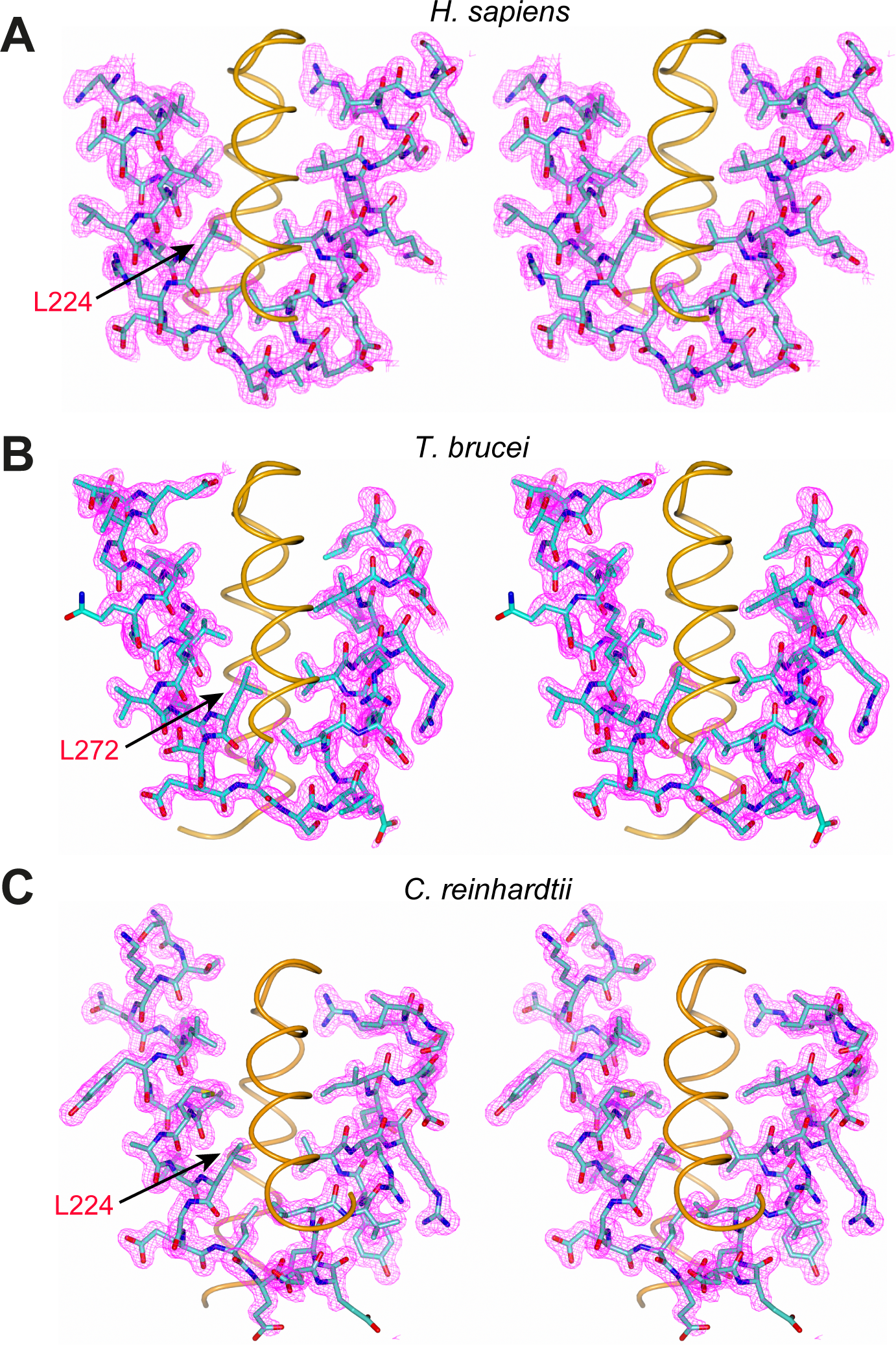
Electron density maps of the CFAP410-CTD crystal structures. (**A**-**C**) Stereo views of the 2*Fo-Fc* maps contoured at 2 σ level for the CTD of *H. sapiens*, *T. brucei* and *C. reinhardtii* CFAP410 proteins. For clarity, only one dimer of the tetramer is shown in each case, with one of the two chains displayed as sticks with electron densities in magenta and the other chain as an orange tube. The conserved disease-causing Leu residues are marked. The plots were produced using CCP4mg.

**Table 1.** Data collection and refinement statistics.

|  | **HsCFAP410-CTD** | **TbCFAP410-CTD** | **CrCFAP410-CTD** |
| --- | --- | --- | --- |
| Wavelength (Å) | 0.873 | 0.972 | 0.972 |
| Resolution range (Å) | 19.94 - 1.50<br>(1.55 - 1.50) | 19.10 - 1.29<br>(1.34 - 1.29) | 19.56 - 1.40<br>(1.45 - 1.40) |
| Space group | C 1 2 1 | P 3<sub>2</sub> 2 1 | P 2<sub>1</sub> 2<sub>1</sub> 2<sub>1</sub> |
| Unit cell<br>*a*, *b*, *c* (Å)<br>$\alpha$, $\beta$, $\gamma$ (°) | 56.12, 50.63, 22.15<br>90, 101.08, 90 | 40.11, 40.11, 62.68<br>90, 90, 120 | 43.44, 48.10, 53.16<br>90, 90, 90 |
| Total reflections | 60,771 (5,760) | 141,666 (12,689) | 275,741 (19,011) |
| Unique reflections | 9,761 (948) | 15,025 (1,430) | 22,140 (1,924) |
| Multiplicity | 6.2 (6.1) | 9.4 (8.9) | 12.5 (9.9) |
| Completeness (%) | 99.60 (98.13) | 99.65 (97.15) | 98.19 (87.57) |
| Mean I/sigma(I) | 11.05 (0.99) | 8.16 (0.76) | 9.40 (1.11) |
| R-merge | 0.077 (1.477) | 0.132 (2.047) | 0.136 (1.019) |
| R-meas | 0.084 (1.616) | 0.140 (2.173) | 0.142 (1.074) |
| R-pim | 0.0335 (0.646) | 0.046 (0.717) | 0.040 (0.331) |
| CC<sub>1/2</sub> | 0.999 (0.519) | 0.991 (0.43) | 0.995 (0.747) |
| CC* | 1.00 (0.827) | 0.998 (0.775) | 0.999 (0.925) |
| Reflections used in refinement | 9,750 (946) | 15,021 (1,430) | 22,131 (1,924) |
| Reflections used for R-free | 977 (95) | 1,516 (141) | 1,988 (173) |
| R-work | 0.213 | 0.187 | 0.165 |
| R-free | 0.249 | 0.204 | 0.190 |
| Number of heavy atoms in proteins | 452 | 505 | 1,084 |
| Number of solvent | 28 | 33 | 91 |
| RMS (bonds) | 0.006 | 0.005 | 0.009 |
| RMS (angles) | 0.93 | 0.62 | 0.93 |
| Ramachandran favored (%) | 100.00 | 100.00 | 98.40 |
| allowed (%) | 0.00 | 0.00 | 1.60 |
| outliers (%) | 0.00 | 0.00 | 0.00 |

Statistics for the highest-resolution shell are shown in parentheses.

All these three crystal structures show a similar homotetrameric arrangement, with each monomer consisting of two α helices that form an open nutcracker-like conformation (**Figure 2A-C**). Despite some variations in the tetramers (**Figure 2D**), all monomers in the three proteins essentially adopt an identical conformation, with root-mean-square deviation (RMSD) of <0.5 Å for all aligned backbone atoms (**Figure 2E**). Each of the two helices consists of approximately 15 residues (**Figure 2F**).

**Figure 2.**
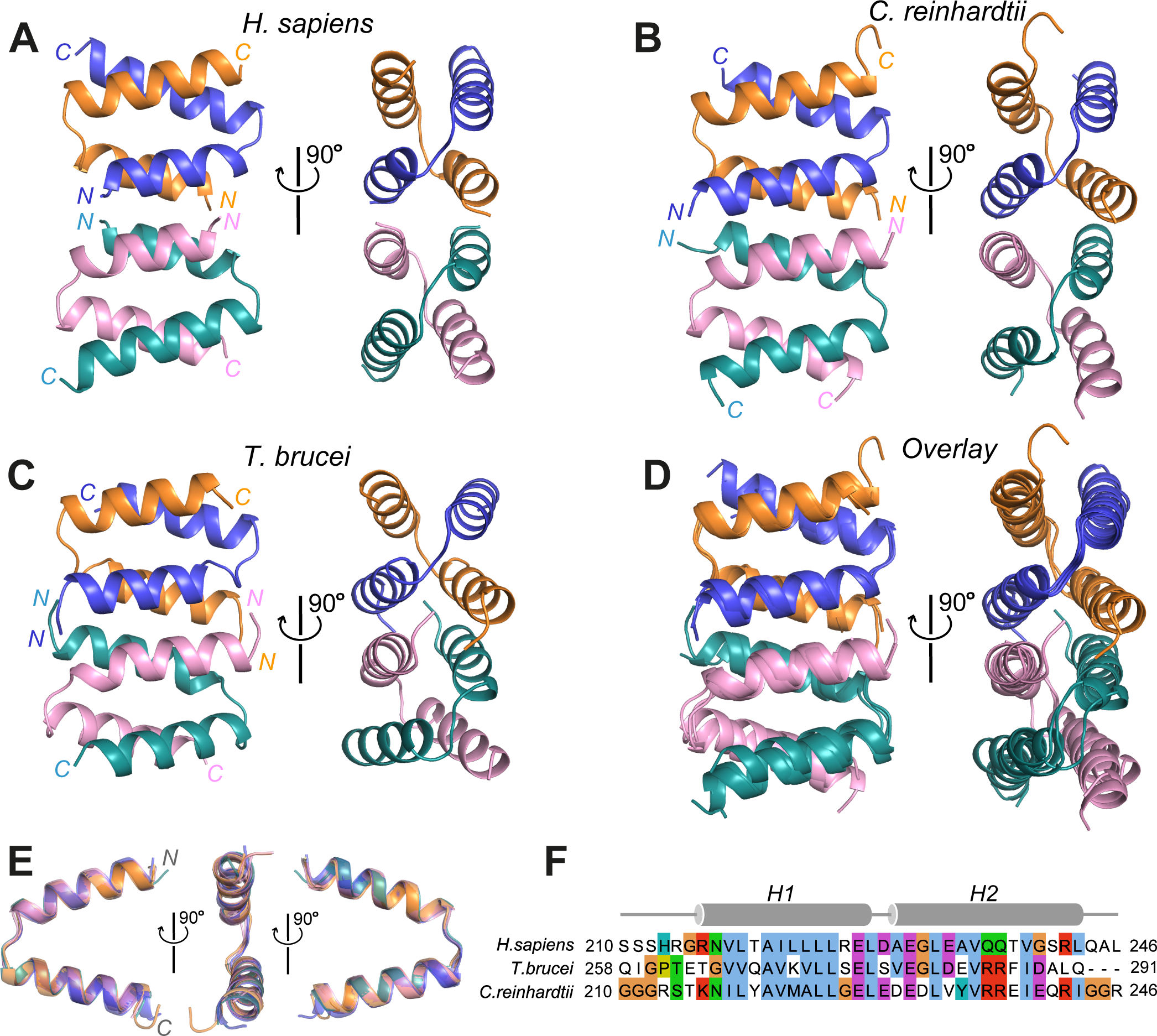
CFAP410-CTD forms a highly conserved tetrameric helical bundle. (**A-C**) Crystal structures of the CTDs of *H. sapiens*, *C. reinhardtii* and *T. brucei* CFAP410. Two orthogonal views are shown in each case. The four copies in each structure are colored differently with their N- and C-terminal ends labelled. (**D**) Superposition of the three tetrameric structures shown in A-C. (**E**) Superposition of all unique monomeric structures from above three tetramers. (**F**) Sequence alignment of the CTDs of *H. sapiens*, *C. reinhardtii* and *T. brucei* CFAP410 with residue ranges shown at both ends. The two α-helices of the crystal structures are shown above the alignments. The alignments were carried out using the option of “Tcoffee with defaults” in Jalview, with residues highlighted using the Clustal color scheme.

### The CFAP410-CTD is a dimer of two individually folded dimers

Multiple single-residue mutations of HsCFAP410 have been identified in patients with SMDAX or RDMS, one of which is L224P located in its CTD [4]. With the crystal structures we were able to examine how this mutation may affect the folding and/or assembly of the domain. We observed in our crystal structures that two CTD monomers form a criss-cross dimer. Two such dimers further pack into a tetramer in a head-to-head manner. The centrally packed N-terminal helix (i.e. H1) consists of mostly hydrophobic residues that are involved in either intra- or inter-dimer interactions, whereas the peripheral C-terminal helix (i.e. H2) is less hydrophobic and contains multiple polar and charged residues (**Figure 3A**). Besides the aforementioned mutation site L224 in H1, there is another highly conserved alanine (Ala) residue that is located in the center of the same helix as L224 (**Figure 3B**). While the Leu residue is fully buried in the dimeric interface (**Figure 3C**), the Ala residue is situated on the hydrophobic inter-dimer interface, with the four equivalent copies arranged in a diamond-like shape across both interfaces of the two dimers (**Figure 3D**; **Figure S3**).

**Figure 3.**
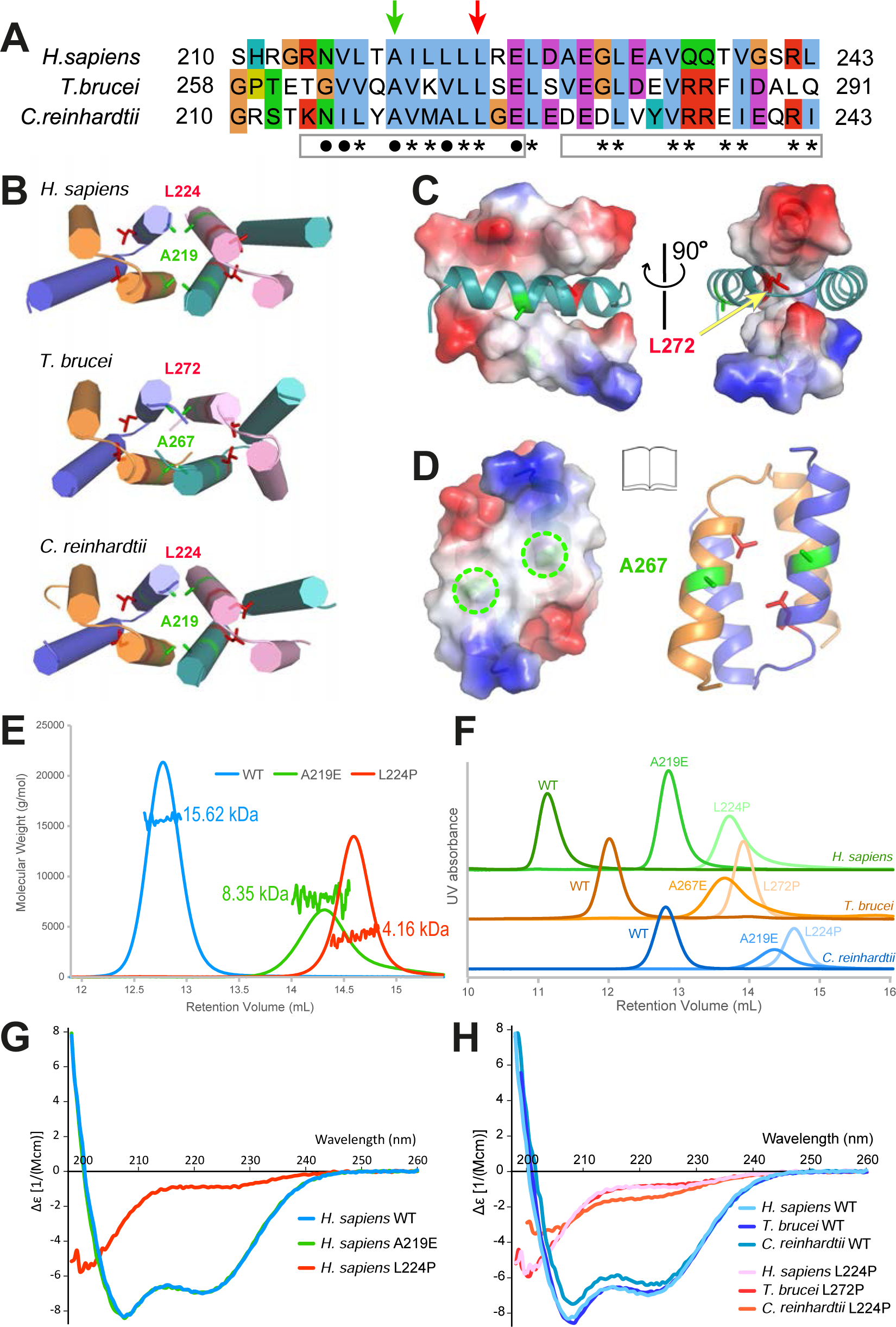
Mutational analyses of CFAP410-CTDs. (**A**) Sequence alignment of the CTDs of three CFAP410 proteins. Boxes under the alignments represent the two α helices in the crystal structure. Residues involved in intra- and inter-dimer interactions are denoted by asterisks and dots, respectively. Green arrow indicates the conserved alanine located at the center of the dimer-dimer interface; red arrow indicates the conserved disease-causing Leu residue. (**B**) Crystal structures of the three CFAP410-CTDs. All helices are depicted as cylinders with the conserved Ala and Leu residues shown as sticks. (**C**) Two orthogonal views of the TbCFAP410-CTD dimer with one chain shown as an electrostatic surface plot and the other as ribbons. The conserved Leu residue (L272) is deeply buried between the two chains. (**D**) An open-book view of the TbCFAP410-CTD tetramer shown in (B), with the dimer on the left shown as an electrostatic surface plot and the one on the right as ribbons. Marked by green circles (left) and shown as sticks (right) are the four conserved Ala residues (A267) situated at the center of the inter-dimer interface. (**E**) SLS results of wild-type (WT), A219E and L224P of CrCFAP410-CTD. (**F**) SEC profiles of WT and mutants of the CTDs of all three CFAP410 proteins. (**G**) CD spectra of WT, A219E and L224P of HsCFAP410-CTD. (**H**) CD spectra of WT and Leu mutants of the CTDs of all three CFAP410 proteins.

To find out whether and how these highly conserved Leu and Ala residues contribute to the formation of the tetramer, we first examined the wild-type and mutants A219E and L224P of CrCFAP410-CTD using the static light scattering (SLS) method (**Figure 3E**). Our data demonstrated that the wild-type protein formed a tetramer as revealed in the crystal structure, but mutant A219E became a dimer. Surprisingly, mutant L224P completely disrupted the oligomeric structure and formed only a monomer. The same results were also observed for the CTDs of *H. sapiens* and *T. brucei* CFAP410 (**Figure 3F**).

We next checked whether either of these two mutants affects the folding of the proteins using the circular dichroism (CD) method. Our results showed that for HsCFAP410-CTD both the wild-type and mutant A219E adopted the same conformation (**Figure 3G**). However, the disease-causing mutant L224P lost all characteristic helical features and became completely unfolded. Consistent results were also seen for both *T. brucei* and *C. reinhardtii* CFAP410 proteins (**Figure 3H**).

To better understand the dramatic influence of the disease-causing mutant L224P on the structural stability of HsCFAP410, we further carried out molecular dynamics simulation analyses on both the wild-type and the mutant. Our conformational analyses of the simulation data showed that while the wild-type model of HsCFAP410-CTD agreed very well with the crystal structure (RMSD = 0.8 Å), the simulated model of L224P was drastically different from the wild-type crystal structure, with the associated all-atom RMSD of 4.4 Å (**Figure 4A**). Such structural changes were seen in all four subunits of the complex structure (**Figure 4B**). The most dramatic difference can be attributed to the major disruption of intra-dimeric contacts (**Figure 4C**). The simulation analyses also indicated significant changes in the hydrophilic and hydrophobic components in mutant L224P in comparison with the wild-type structure (**Figure 4D, E**).

**Figure 4.**
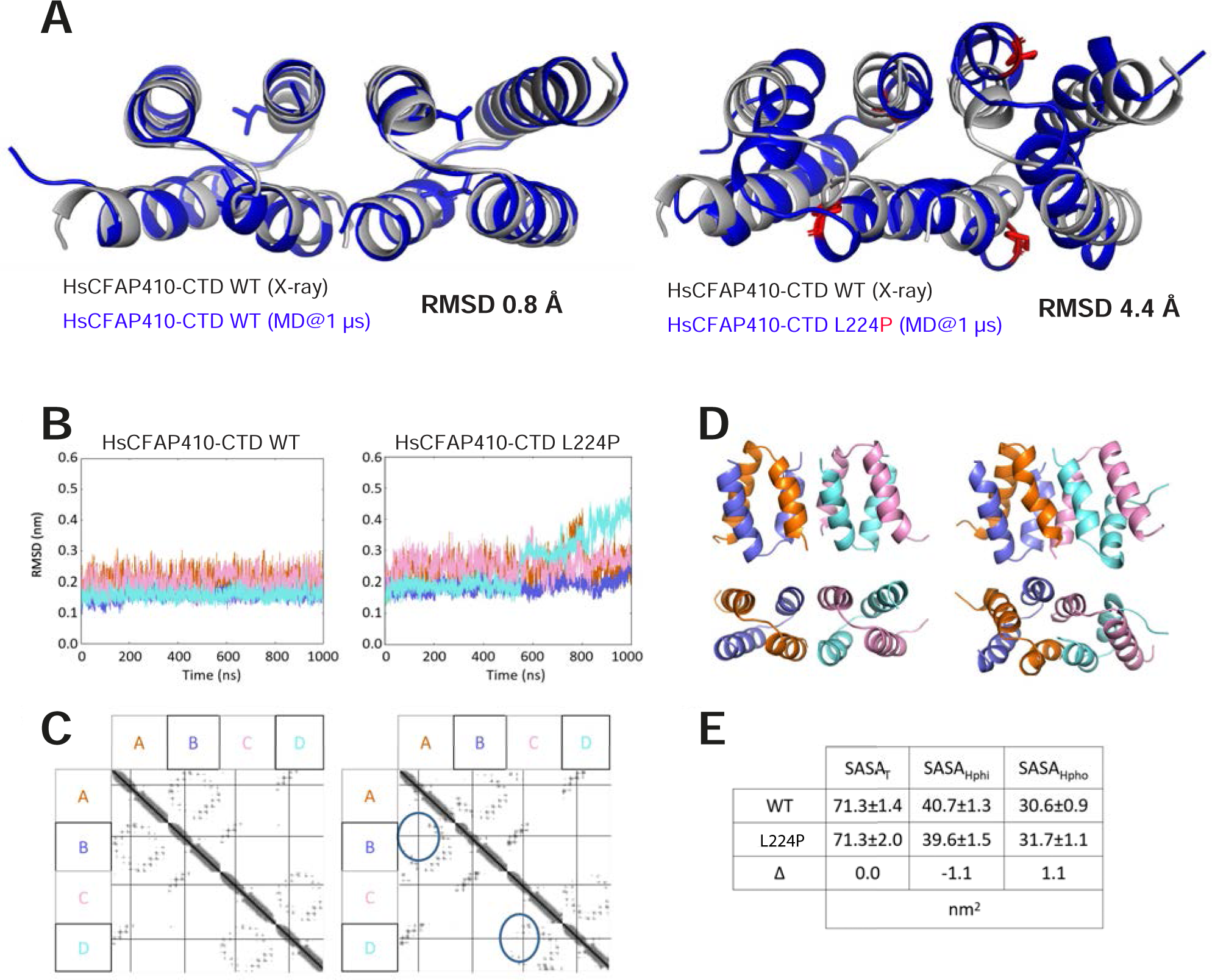
Simulation analyses of wild type and the L223P mutant of HsCFAP410-CTD. (**A**) Superposition of the crystal structure of the wild-type (WT) HsCFP0-CTD with the simulated WT (left) or the simulated L224P mutant (right) after 1 µs of molecular dynamics, with the associated all-atom RMSD indicated. (**B**) Comparison of all-atom RMSD from the initial structure for hCTD WT and L224P simulations across different domains, as indicated by color. (**C**) Contact maps from WT (left) and L224P (right) simulations with the major disruption of contacts between chains A and B as well as C and D, as induced by the L224P mutations, indicated by blue ovals. (**D**) The 1 ms snapshots of WT (left) and L224P (right) HsCFP0-CTD colored according to the scheme given in (**C**). (**E**) Comparison of the total solvent-accessible surface area (SASA_T_) and its hydrophilic (SASA_Hphi_) and hydrophobic components (SASA_Hpho_) in WT and L224P mutant with the change indicated by Δ (Δ = SASA_mutant_-SASA_wt_).

### TbCFAP410 is essential for cytokinesis

We next examined the function of CFAP410 *in vivo*. First of all, we examined the localization of TbCFAP410 in *T. brucei* by endogenously tagging the protein with an N-terminal mNeonGreen fluorescent protein. The fused protein was expressed in a cell line that also expresses SAS6, a basal body marker, endogenously tagged with mScarlet. We observed that in 1K1N cells, mNG::CFAP410 was concentrated at the posterior cell tip, with a slight enrichment observed at the basal body region above the cytoplasmic signal (**Figure 5A**, left). As the cell cycle progressed to the 2K1N and 2K2N stages, the mNG::CFAP410 signal extended as a line from the posterior cell tip along the ventral edge of the cell. We further generated detergent-extracted cytoskeletons to confirm whether mNG::CFAP410 was stably associated with the cytoskeleton (**Figure 5A**, right). In cytoskeletons of 1K1N cells, mNG::CFAP410 was concentrated at the posterior cell tip, with an additional weaker signal from the basal body region. Later in the cell cycle (2K1N and 2K2N), the mNG::CFAP410 signal extended from the posterior cell tip along the ventral edge, with an additional signal from the now duplicated basal body region. This suggests that CFAP410 is associated with microtubule organizing centers in *T. brucei*, predominantly the posterior cell tip.

**Figure 5.**
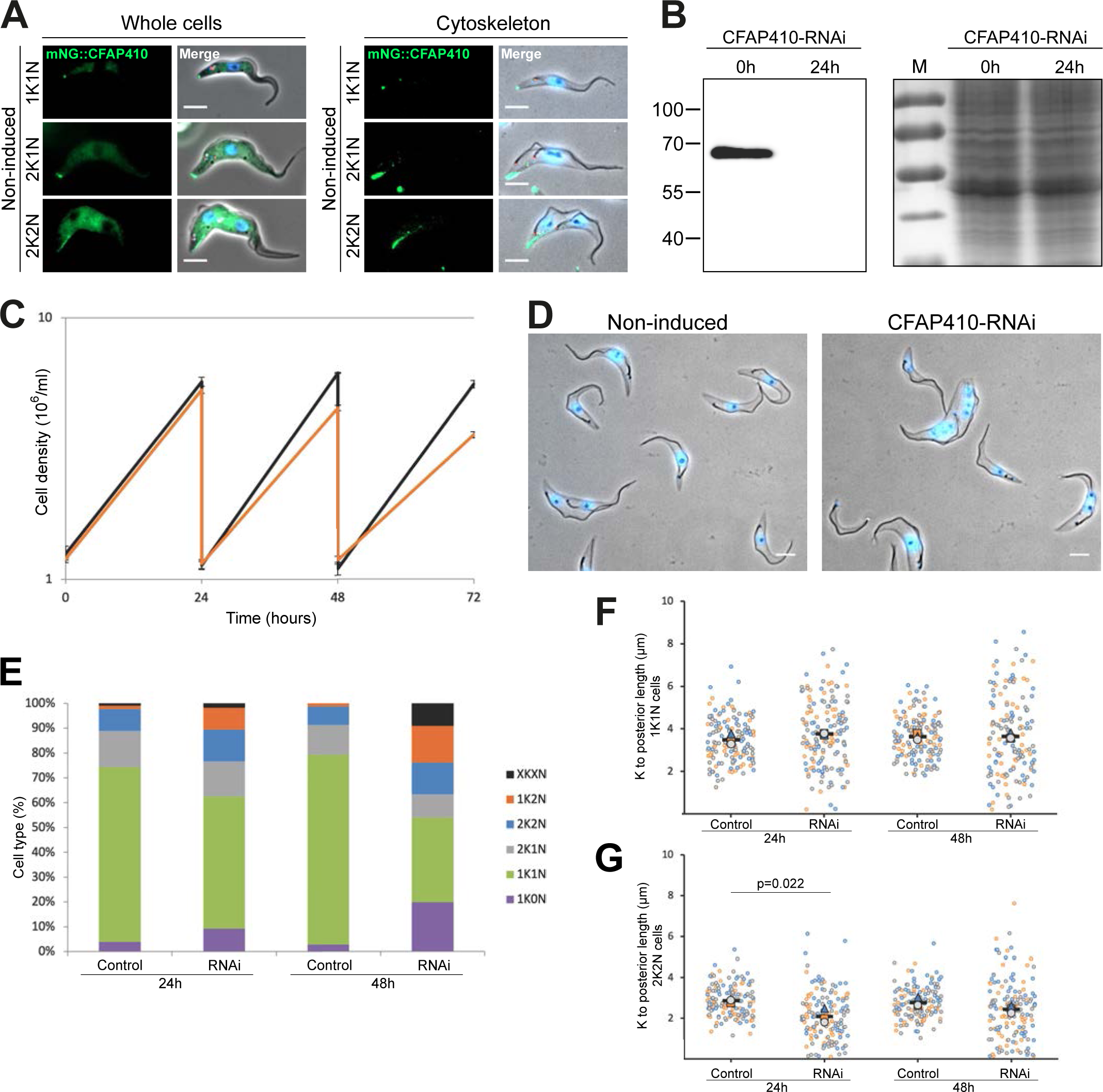
TbCFAP410 depletion caused a cytokinesis defect in *T. brucei*. (**A**) Fluorescence microscopy images of live cells or methanol fixed cytoskeletons expressing mNG::TbCFAP410 (green) and mSc::TbSAS6 - basal body marker (red). Scale bar: 5 μm. *K: kinetoplast. N: nuclei*. (**B**) Western blot (left) and Coomassie stained gel (right) showing induction of TbCFAP410-RNAi for 24 h in *T. brucei*. BB2 (anti-Ty) antibody was used. 4×10^6^ parasites per lane were loaded. *M: marker.* (**C**) Growth curve of non-induced (black) and CFAP410-RNAi induced (orange) cells. Induction was repeated three times, and standard deviation is shown. (**D**) Field of view of detergent extracted cytoskeletons of non-induced cells and CFAP-410 RNAi cells induced for 48 hours. Scale bar: 5 μm. (**E**) Quantitation of cell types seen in non-induced (control) and TbCFAP410-RNAi. Percentages were calculated from number of cell categories seen after cytoskeleton extraction from ≥500 cells. Induction was repeated three times, with a similar result each time; data from one is shown. *XKXN: aberrant number of kinetoplast and nuclei.* (**F, G**) Distance between kinetoplast and the posterior cell tip in 1K1N (**F**) and 2K2N (**G**) in cytoskeletons of non-induced (control) and induced TbCFAP410-RNAi. Each dot represents the length measurement of an individual cytoskeleton, and the colors represent three replicates. The larger triangle, circle and square are the average of each replicate, and the black bar is the average between the three replicates. For each replicate, 50 cells were measured. p-values were calculated using independent samples t-test.

Next, we depleted TbCFAP410 by RNAi to determine its function in *T. brucei*. Depletion of CFAP410 after RNAi was confirmed by western blot, with a complete loss of mNG::TbCFAP410 protein after 24 h of induction (**Figure 5B**). We observed that TbCFAP410 depletion substantially reduced cell growth by 48 h after induction (**Figure 5C**). To understand the causes of the growth defect, we further quantified the cell types present in the culture after RNAi-induction. After 24 h, the depletion of TbCFAP410 caused a minor reduction in the number of 1K1N cells, with a concomitant increase in 1K0N (zoids) and 1K2N cells, when compared to the non-induced groups (**Figure 5D, E**). This effect continued with more aberrant cells, with multiple kinetoplasts and nuclei appearing at 48 h. This suggests that TbCFAP410 plays an important role in *T. brucei* cytokinesis.

After TbCFAP410 RNAi induction, we noted that there were many cells in which the kinetoplast was very close to the posterior cell tip and other cells with an elongated posterior cell tip. To investigate this further, we measured the kinetoplast to the posterior cell tip in detergent-extracted cytoskeletons. Although we observed only little change in the average distance from the posterior cell tip to kinetoplast in 1K1N cytoskeletons after induction, there was a substantial increase in the range of these measurements, with cytoskeletons observed having a more reduced or increased distance from the kinetoplast to the posterior cell tip (**Figure 5F**). Moreover, in 2K2N cells the posterior kinetoplast was positioned significantly closer to the posterior cell tip after RNAi induction (**Figure 5G**). These data suggest that TbCFAP410 is required for the regulation of the posterior cell tip elongation during the cell cycle.

### The tetrameric assembly of the CFAP410-CTD is essential for its localization

We next examined which part of CFAP410 controls its specific localization to the basal body and how the disease-causing mutation L224P affects its function *in vivo*. All target proteins were expressed in *T. brucei* using a tetracycline-inducible expression system, with an N-terminal mNeonGreen fluorescent protein tag. Since the cell lines had a strong cytoplasmic signal, we also generated detergent-extracted cytoskeletons to categorize the localization of the TbCFAP410 mutants (**Figure 6A**). The mNG-tagged wild-type protein localized to both the posterior end and the basal body region, as seen with the endogenously tagged protein. In the mutant A267E, the mNG::CFAP410 signal was exclusively found at the posterior of most cytoskeletons (63.5%), while full-length TbCFAP410-L272P and the two CTD-lacking constructs, NTD and NTD-linker, did not show any signal in cytoskeletons (**Figure 6B**). Notably, all constructs were induced for 24 h and their expression was confirmed by western blot (**Figure 6C, D**). This suggests that both the presence of the CTD and the integrity of its oligomerization are essential for the interaction of TbCFAP410 with the basal body and posterior cell tip.

**Figure 6.**
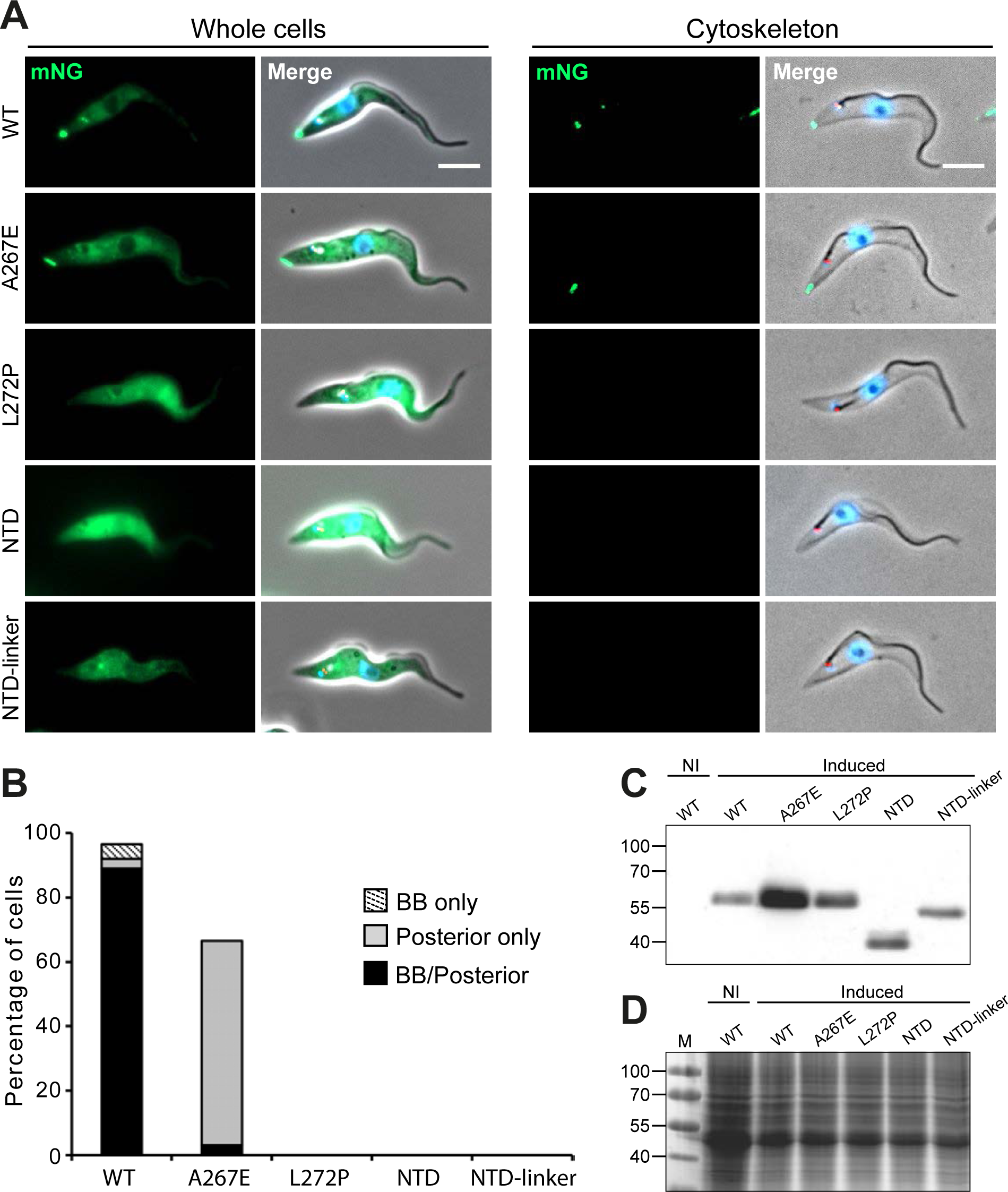
TbCFAP410 C-terminal mutants have a disrupted localization. (**A**) Images acquired from live cells or methanol fixed cytoskeletons from doxycycline induced cells expressing TbCFAP410 wild-type (WT) or TbCFAP410 mutants (green) with mSc::TbSAS6 - basal body marker (red). Scale bar: 5 μm. (**B**) Histogram of TbCFAP410 signal counts. The percentage of cytoskeletons with a basal body and posterior signal, a basal body only signal and posterior signal only was calculated for each cell line. Induction was repeated twice, with a similar result both times and data from one is shown. *BB: basal body.* (**C, D**) Western blot (**C**) and Coomassie stained gel (**D**) showing the induction of WT and truncated or mutant TbCFAP410 expression in *T. brucei*. BB2 (anti-Ty) antibody was used. 4×10^6^ parasites per lane were loaded. *NI: non-induced*.

## Discussion

CFAP410 is a protein present in all ciliates and plays an essential role in ciliogenesis [4]. Located at the basal body, the protein is highly conserved across phyla including algae, protista and animals. Here we report our structural studies on the CTD of three orthologs of CFAP410 from *T. brucei*, *C. reinhardtii*, and *H. sapiens*. CFAP410 is a bimodular protein comprising two folded domains, i.e. NTD and CTD, which are connected via a variable unstructured loop (**Figure S1**). The crystal structures show that all three CTDs adopt a similar nutcracker-like conformation composed of two helices (**Figure 1**). Two CTDs first form an intercalated homodimer, and two such dimers further assemble into a tetramer by packing head-to-head via their first helices (**Figure 2**).

Multiple single-residue mutations of HsCFAP410 have been identified in patients with SMS or RDMS ciliopathies [13-15]. One of these mutations, L224P, is located in the CTD. Our biochemical studies showed that such mutation not only disrupts its helical property but also causes complete disassembly of the tetramer (**Figure 3**). Consistently, our simulation analyses indicated significant changes in the hydrophilic and hydrophobic components in mutant L224P in comparison with the wild-type structure (**Figure 4**). Our results also suggest that CFAP410-CTD consists of two independently folded dimers, as mutation of the highly conserved Ala residue on the inter-dimer interface disassembles the tetramer to two dimers, but the dimer maintains the same secondary structure as the wild-type protein.

Our *in vivo* data showed that TbCFAP410 localizes to both the basal body and the posterior cell tip. Depletion of TbCFAP410 by RNAi not only caused growth defects, but also resulted in aberrant cells with multiple kinetoplasts and nuclei appearing, demonstrating that TbCFAP410 plays an important function in cytokinesis (**Figure 5**). We observed that the A267E mutant that breaks the tetramer into two dimers could still localize to the posterior cell tip, but failed to target to the basal body. However, the equivalent disease-causing mutant L272P of TbCFAP410 completely lost its localization to both the posterior cell tip and the basal body (**Figure 6**). It suggests that targeting TbCFAP410 to the basal body requires both the tetrameric assembly and the correct folding of its CTD. Consistently, we found that two truncated constructs containing either the NTD alone or the NTD together with the linker failed to localize to either the basal body or posterior cell tip.

Previous immunofluorescence studies showed that CFAP410 co-localizes with both the serine/threonine kinase NEK1 and a retinal ciliopathy protein called SPATA7 at the basal body in hTERT-RPE1 cells [4]. Other studies also showed that CFAP410 is a component of a retinal ciliopathy-associated protein complex containing both NEK1 and SPATA7 [16]. Direct robust interaction between CFAP410 and NEK1 has recently been observed in human cells [17]. The interaction possibly involves both the NTD and the CTD of CFAP410. It was shown previously that the L224P mutant of CFAP410 abolishes its interaction with NEK1 [4]. Given that the mutant L224P not only disassembles the tetramer of CFAP410-CTD but also causes the unfolding of the helical structure (**Figure 3**), the tetrameric assembly of CFAP410 seems to play an essential role in its interaction with NEK1. Therefore, the disrupted location of TbCFAP410-L272P to the basal body we observed here could be attributed to its abolished interaction with NEK1 as what occurs in human cells.

In summary, taking together our structural, biochemical and *in vivo* studies, we conclude that CFAP410 forms a tetramer via its CTD, which fuses the four globular NTDs together to form an oligomeric protein complex (**Figure 7**). The tightly packed eight-helix bundle of the CTD controls the specific localization of CFAP410 to the basal body possibly by directly interacting with the serine/threonine kinase NEK1. The single-residue mutation L224P in HsCFAP410 that causes the autosomal recessive ciliopathy SMDAX could thus be attributed to the disruption of its oligomeric assembly that abolishes its interaction with a partner protein at the basal body.

**Figure 7.**
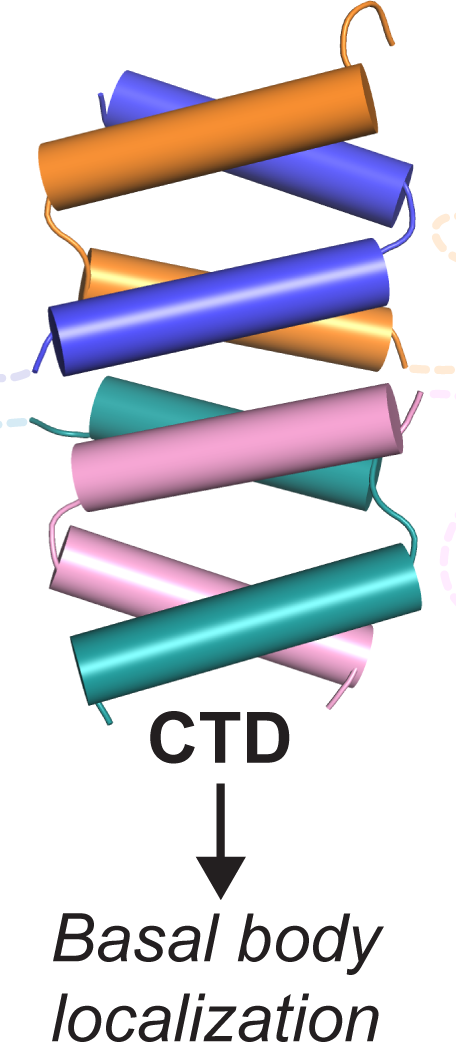
Assembly and basal body localization control of CFAP410. CFAP410-CTD forms a tetrameric helical bundle, which is connected to the globular NTD via a long disordered loop. The tetrameric CTD controls the specific localization of CFAP410 to the basal body.

## Materials and Methods

### Preparation of synthetic polypeptides

Wild-type and mutant polypeptides of the C-terminal region of *T. brucei, H. sapiens* and *C. reinhardtii* CFAP410 were chemically synthesized and purified by an in-house facility. All HsCFAP410-CTD fragments were soluble in a buffer containing 20 mM HEPES pH 7.5, 100 mM NaCl, and 1 mM DTT. However, those of TbCFAP410 and CrCFAP410 only partially dissolved in the buffer. We therefore carried out refolding for the insoluble polypeptides using the following protocol. Dry peptide powder was first dissolved in 8 M urea at a concentration of 10 mg/ml, and the solution was then quickly diluted by 10 × fold using the above buffer. After dialysis against the same buffer overnight at 4°C, the samples were loaded onto a Superdex 75 Increase 10/300 GL column (Cytiva). Fractions containing the target polypeptides were pooled and concentrated to 10-15 mg/ml. Sample homogeneity was confirmed on both SDS PAGE and native polyacrylamide gels.

### Crystallization and structural determination

Purified synthetic CTDs of all CFAP410 proteins were subjected to extensive crystallization trials using multiple commercial crystallization kits (Hampton Research). Conditions of the initial hits were further optimized to obtain single crystals. The final condition for crystallizing the TbCFAP410-CTD contained 2.0 M (NH_4_)_2_SO_4_ and 5% (v/v) iso-propanol (4°C). The crystallizing condition for HsCFAP410-CTD contained 1.5 M NaCl and 10% (v/v) ethanol (22°C). CrCFAP410-CTD was crystallized in a condition containing 0.02 M CaCl_2_, 0.1 M sodium acetate (pH 5.0) and 30% (v/v) 2-Methyl-2,4-pentanediol at 4°C.

All X-ray diffraction data were collected at the ESRF synchrotron site and processed using XDS [18]. The crystal structure of TbCFAP410-CTD was determined *de novo* using ARCIMBOLDO [19]. Those of HsCFAP410-CTD and CrCFAP410-CTD were determined subsequently by molecular replacement using the crystal structure of TbCFAP410-CTD as a search template. All resulting models were optimized by multiple rounds of manual rebuilding in COOT [20] and refinement in Phenix [21], and finally validated in MolProbity [22]. Details of data collection and refinement are summarized in **Table 1**.

### Static light scattering (SLS)

SLS measurements were carried out by coupling SEC with mass determination. 50 μl of purified polypeptides at 2 mg/ml was analyzed on a Superdex S-200 10/300 GL column (Cytiva) pre-equilibrated with a buffer containing 20 mM HEPES (pH 7.5), 100 mM NaCl, 1 mM DTT, and 1% (v/v) glycerol. The polypeptides were run at a flow rate of 0.5 ml/min on a high-performance liquid chromatography (HPLC) system (Agilent Technologies 1260 infinity) which was directly connected to a Mini-DAWN Treos light-scattering instrument (Wyatt Technology Corp., Santa Barbara, CA). Data analyses were carried out using ASTRA software provided by the manufacturer. Final curves were built in SigmaPlot [23].

### Circular dichroism (CD)

Far-UV CD spectra of all polypeptides between 195 and 260 nm were measured on a Chirascan plus CD spectrometer (Applied Photophysics) in a cuvette with a 0.5-mm path length. Samples were diluted to approximately 0.2 mg/ml using a buffer containing 10 mM HEPES (pH 7.5) and 100 mM NaF. Data points were corrected for buffer signals and drifts. CD curves were generated using SigmaPlot by averaging data collected from two scans for each protein sample.

### Generation of constructs for *in vivo* assays and cell culture of *T. brucei*

The DNA sequence of TbCFAP410 was recoded in wild-type and all mutations for *in vivo* assays. The recoded DNA sequences were chemically synthesized (Twist Bioscience) and cloned into the plasmid pDEX577-mNG. All constructs were subsequently linearized with *Not*I and then electroporated using a BTX Gemini Twin Wave with 3 pulses of 1.7 kV for 100 µs as previously described [24]. Cells were recovered in SDM-79 for 8 hours before selection with phleomycin (5 µg/ml).

SmOxP9 procyclic *T. brucei* expressing TbSAS6 endogenously tagged with mScarlet were used for all experiments [25]. These were derived from the TREU 927 strain, expressing T7 RNA polymerase and tetracycline repressor [26] and were grown at 28°C in SDM-79 medium supplemented with 10% FCS [27]. Cell concentration was determined in a Z2 Coulter Counter particle counter.

### Fluorescence microscopy

Cells were induced overnight with doxycycline (0.5 µg/ml) harvested by centrifugation at 800×g for 5 min. For live-cell microscopy, cells were washed two times in PBS supplemented with 10 mM glucose and 46 mM sucrose (vPBS). In the last wash, the DNA was stained using 10 µg/ml Hoechst 33342 (Sigma-Aldrich), re-suspended in vPBS and then mounted with a coverslip and immediately imaged. For cytoskeletons, cells were washed in PBS, settled onto glass slides for 5 min and treated with 1% (v/v) NP-40 in PEME for 1 min. Cytoskeletons were then fixed in −20°C methanol for 20 min and rehydrated in PBS for 10 min. After fixation, the DNA was stained with 20 μg/ml Hoechst 33342 in PBS, washed in PBS for 10 min or overnight and mounted before imaging. Images were taken using a Zeiss Axio Imager.Z1 microscope equipped with an ORCA-Flash 4.0 CMOS camera using a Plan-Apochromat 63×/1.4 NA oil objective. Images were acquired and analyzed with ZEN 2 PRO software and assembled for publication in Adobe Illustration CS6.

### Molecular dynamics (MD) simulations

DSF measurements were performed on a CFX Connect Real-Time PCR machine (Bio-rad). Proteins were diluted to 0.4 mg/ml using a buffer containing 10 mM Tris-HCl pH 8.0 and 100 mM NaCl. To prepare for the measurements, 24 μl of the protein sample was mixed with 1 μl of 20× SYBR orange dye (ThermoFisher Scientific). Each sample was prepared and measured in duplicates, and the two measurements were averaged to produce the final curve. Melting temperature (T_m_) values were obtained using the software provided by the manufacturer.

Atomistic models for wild-type TbCFAP410-NTD and all mutants (V37F, C63Y, R75P, Y107C, Y107H and V111M) were constructed starting from the crystal structure by *in silico* mutagenesis using Pymol (http://www.pymol.org). The same strategy was followed with the wild-type and single-residue mutants of the tetrameric CTD structures. All further calculations were performed using the GROMACS 2019 package [28] and Amber99SB-ILDN force-field parameters [29]. In all cases, the simulated system was placed into a 7×7×7-nm^3^ octahedral box, energy minimized and solvated with TIP3P water molecules. Additionally, Na^+^ and Cl^−^ ions were added to achieve electroneutrality at the final salt concentration of 150 mM. The complete system was then energy-minimized using position restraints placed on protein Cα-atoms, and subjected to 5,000 steps of NVT molecular dynamics (MD) with a 0.5-fs time step and subsequent 250,000 steps of NPT MD with a 1-fs time step. After this initial equilibration, a 1 μs NPT production MD run was simulated (5×10^8^ steps with a 2-fs time step) for each system. All analysis was performed on the final 750 ns of each simulated trajectory. A twin-range (10 Å, 10.5 Å) spherical cut-off function was used to truncate the van der Waals interactions. Electrostatic interactions were treated using the particle-mesh Ewald summation (real space cut-off of 10-Å and 1.2-Å grid with fourth-order spline interpolation). MD simulations were carried out using 3D periodic boundary conditions, in the isothermal−isobaric (NPT) ensemble with an isotropic pressure of 1.013 bar and a constant temperature of 295 K. The pressure and temperature were controlled using a Nose-Hoover thermostat [30, 31] and a Parrinello–Rahman barostat with 0.5-ps and 10-ps relaxation parameters, respectively, and a compressibility of 4.5×10^−5^ bar^−1^ for the barostat.

### Accession code

Coordinates and structure factors have been deposited in the Protein Data Bank (PDB) under accession codes 8AXO, 8AXQ, and 8AXR.

## Acknowledgments

We are grateful to the staff at the beamlines ID23-1 and ID23-2 at the European Synchrotron Radiation Facility (ESRF) for their help with X-ray diffraction data collection. We also thank Dr. A. Burga for helping us order all synthetic polypeptides. This work was supported by the grant I5960-B2 from the Austrian Science Fund (FWF) to GD and a Newton International Fellowship (NIF\R1\191618) to HG. LDL was supported by a CAPES fellowship and a Company of Biologists Travel Award. Work in the JS lab is supported by the Wellcome Trust and Leverhulme Trust. We acknowledge the support of the Oxford Brookes Centre for Bioimaging. SA acknowledges the VIP2 fellowship 2020 funded by the European Union’s Horizon 2020 research and innovation programme under the Marie Skłodowska-Curie grant agreement No. 847548.

## References

1. Krohn, K., et al., Immunochemical characterization of a novel mitochondrially located protein encoded by a nuclear gene within the DFNB8/10 critical region on 21q22.3. Biochem Biophys Res Commun, 1997. 238(3): p. 806–10.

2. Abu-Safieh, L., et al., Autozygome-guided exome sequencing in retinal dystrophy patients reveals pathogenetic mutations and novel candidate disease genes. Genome Res, 2013. 23(2): p. 236–47.

3. Scott, H.S., et al., Characterization of a novel gene, C21orf2, on human chromosome 21q22.3 and its exclusion as the APECED gene by mutation analysis. Genomics, 1998. 47(1): p. 64–70.

4. Wheway, G., et al., An siRNA-based functional genomics screen for the identification of regulators of ciliogenesis and ciliopathy genes. Nat Cell Biol, 2015. 17(8): p. 1074–1087.

5. Wang, Z., et al., Axial spondylometaphyseal dysplasia is also caused by NEK1 mutations. J Hum Genet, 2017. 62(4): p. 503–506.

6. Fang, X., et al., The NEK1 interactor, C21ORF2, is required for efficient DNA damage repair. Acta Biochim Biophys Sin (Shanghai), 2015. 47(10): p. 834–41.

7. Chia, R., A. Chio, and B.J. Traynor, Novel genes associated with amyotrophic lateral sclerosis: diagnostic and clinical implications. Lancet Neurol, 2018. 17(1): p. 94–102.

8. Shim, K.S., et al., Reduction of chromatin assembly factor 1 p60 and C21orf2 protein, encoded on chromosome 21, in Down syndrome brain. J Neural Transm Suppl, 2003(67): p. 117–28.

9. Ma, X., R. Peterson, and J. Turnbull, Adenylyl cyclase type 3, a marker of primary cilia, is reduced in primary cell culture and in lumbar spinal cord in situ in G93A SOD1 mice. BMC Neurosci, 2011. 12: p. 71.

10. Ehara, S., et al., Axial spondylometaphyseal dysplasia. Eur J Pediatr, 1997. 156(8): p. 627–30.

11. Warman, M.L., et al., Nosology and classification of genetic skeletal disorders: 2010 revision. Am J Med Genet A, 2011. 155A(5): p. 943-68.

12. Wang, Z., et al., Axial Spondylometaphyseal Dysplasia Is Caused by C21orf2 Mutations. PLoS One, 2016. 11(3): p. e0150555.

13. McInerney-Leo, A.M., et al., Homozygous variant in C21orf2 in a case of Jeune syndrome with severe thoracic involvement: Extending the phenotypic spectrum. Am J Med Genet A, 2017. 173(6): p. 1698–1704.

14. Khan, A.O., et al., C21orf2 is mutated in recessive early-onset retinal dystrophy with macular staphyloma and encodes a protein that localises to the photoreceptor primary cilium. Br J Ophthalmol, 2015. 99(12): p. 1725–31.

15. Suga, A., et al., Identification of Novel Mutations in the LRR-Cap Domain of C21orf2 in Japanese Patients With Retinitis Pigmentosa and Cone-Rod Dystrophy. Invest Ophthalmol Vis Sci, 2016. 57(10): p. 4255–63.

16. Eblimit, A., et al., Spata7 is a retinal ciliopathy gene critical for correct RPGRIP1 localization and protein trafficking in the retina. Hum Mol Genet, 2015. 24(6): p. 1584–601.

17. Gregorczyk, M., et al., Functional characterization of C21ORF2 association with the NEK1 kinase mutated in human in diseases. Life Sci Alliance, 2023. 6(7).

18. Kabsch, W., Xds. Acta Crystallogr D Biol Crystallogr, 2010. 66(Pt 2): p. 125–32.

19. Rodriguez, D.D., et al., Crystallographic ab initio protein structure solution below atomic resolution. Nat Methods, 2009. 6(9): p. 651–3.

20. Emsley, P. and K. Cowtan, Coot: model-building tools for molecular graphics. Acta Crystallogr D Biol Crystallogr, 2004. 60(Pt 12 Pt 1): p. 2126-32.

21. Adams, P.D., et al., PHENIX: a comprehensive Python-based system for macromolecular structure solution. Acta Crystallogr D Biol Crystallogr, 2010. 66(Pt 2): p. 213–21.

22. Chen, V.B., et al., MolProbity: all-atom structure validation for macromolecular crystallography. Acta Crystallogr D Biol Crystallogr, 2010. 66(Pt 1): p. 12–21.

23. Kornbrot, D., Statistical software for microcomputers: SigmaPlot 2000 and SigmaStat2. Br J Math Stat Psychol, 2000. 53 (Pt 2): p. 335–7.

24. Dean, S., et al., A toolkit enabling efficient, scalable and reproducible gene tagging in trypanosomatids. Open Biol, 2015. 5(1): p. 140197.

25. Atkins, M., et al., CEP164C regulates flagellum length in stable flagella. J Cell Biol, 2021. 220(1).

26. Poon, S.K., et al., A modular and optimized single marker system for generating Trypanosoma brucei cell lines expressing T7 RNA polymerase and the tetracycline repressor. Open Biol, 2012. 2(2): p. 110037.

27. Brun, R. and Schonenberger, *Cultivation and in vitro cloning or procyclic culture forms of Trypanosoma brucei in a semi-defined medium. Short communication*. Acta Trop, 1979. 36(3): p. 289–92.

28. Van Der Spoel, D., et al., GROMACS: fast, flexible, and free. J Comput Chem, 2005. 26(16): p. 1701–18.

29. Lindorff-Larsen, K., et al., Improved side-chain torsion potentials for the Amber ff99SB protein force field. Proteins, 2010. 78(8): p. 1950–8.

30. Nose, S., A Unified Formulation of the Constant Temperature Molecular-Dynamics Methods. Journal of Chemical Physics, 1984. 81(1): p. 511–519.

31. Hoover, W.G., Canonical Dynamics - Equilibrium Phase-Space Distributions. Physical Review A, 1985. 31(3): p. 1695–1697.

