## Supplemental materials for "The C-terminus of CFAP410 forms a tetrameric helical bundle that is essential for its localization to the basal body"

### Legends for supplemental figures

**Figure S1. Folding prediction and conservation analyses of *H. sapiens* and *T. brucei* CFAP410.** (**A**) Disorder prediction of HsCFAP410 by ODiNPred [1]. The central region spanning residues ~130-210 was predicted to have very high probabilities of being disordered. (**B**) Folding prediction by FoldIndex [2] with a sliding window of 40 suggested an intrinsically unfolded region around residues 125-200 in HsCFAP410. (**C**) Disorder prediction of TbCFAP410 by ODiNPred. The central region spanning residues 200-260 was predicted to have very high probabilities of being disordered. (**D**) Folding prediction by FoldIndex with a sliding window of 40 suggested an intrinsically unfolded region around residues 160-250 in TbCFAP410. (**E**) Sequence alignment of TbCFAP410 homologs from various trypanosome species. The alignment was carried out using the option of "Muscle with defaults" in Jalview [3], with shading indicating the degree of conservation. The predicted unfolded region (residues 161-257) is not conserved and has variable lengths in different species, whereas high conservations were seen in both NTD and CTD. Residue L272 of TbCFAP410 corresponding to the disease-causing mutation L224P is marked.

**Figure S2. Sequence alignments of CFAP410 proteins from various mammalian, kinetoplastid and algae species.** (**A**-**C**) Sequence alignments of close homologs of *H. sapiens*, *T. brucei* and *C. reinhardtii* CFAP410 proteins, respectively. There are two highly conserved domains in each case, which are marked by light blue and orange bars above the alignments. (**D**) Sequence alignment of *H. sapiens*, *T. brucei* and *C. reinhardtii* CFAP410 proteins. The green and red arrows indicate the highly conserved Ala residues on the inter-dimeric interface and the counterparts of the disease-causing L224P mutation in HsCFAP410, respectively.

**Figure S3. Inter-molecular interactions in the HsCFAP410-CTD tetramer.** The plots were generated using DIMPLOT in the LigPlot plus suite [4]. Residues involved in hydrogen bond formation are shown as ball-and-sticks, with oxygen, nitrogen and carbon atoms colored in red, blue, and gray, respectively. Dotted lines indicate hydrogen bonds with inter-atomic distances labeled. Non-bonded residues involved in hydrophobic interactions are shown as spoked arcs.

### References


1. Dass, R., F.A.A. Mulder, and J.T. Nielsen, *ODiNPred: comprehensive prediction of protein order and disorder.* Sci Rep, 2020. **10**(1): p. 14780.

2. Prilusky, J., et al., *FoldIndex: a simple tool to predict whether a given protein sequence is intrinsically unfolded.* Bioinformatics, 2005. **21**(16): p. 3435-8.

3. Waterhouse, A.M., et al., *Jalview Version 2--a multiple sequence alignment editor and analysis workbench.* Bioinformatics, 2009. **25**(9): p. 1189-91.

4. Laskowski, R.A. and M.B. Swindells, *LigPlot+: multiple ligand-protein interaction diagrams for drug discovery.* J Chem Inf Model, 2011. **51**(10): p. 2778-86.

Figure S1

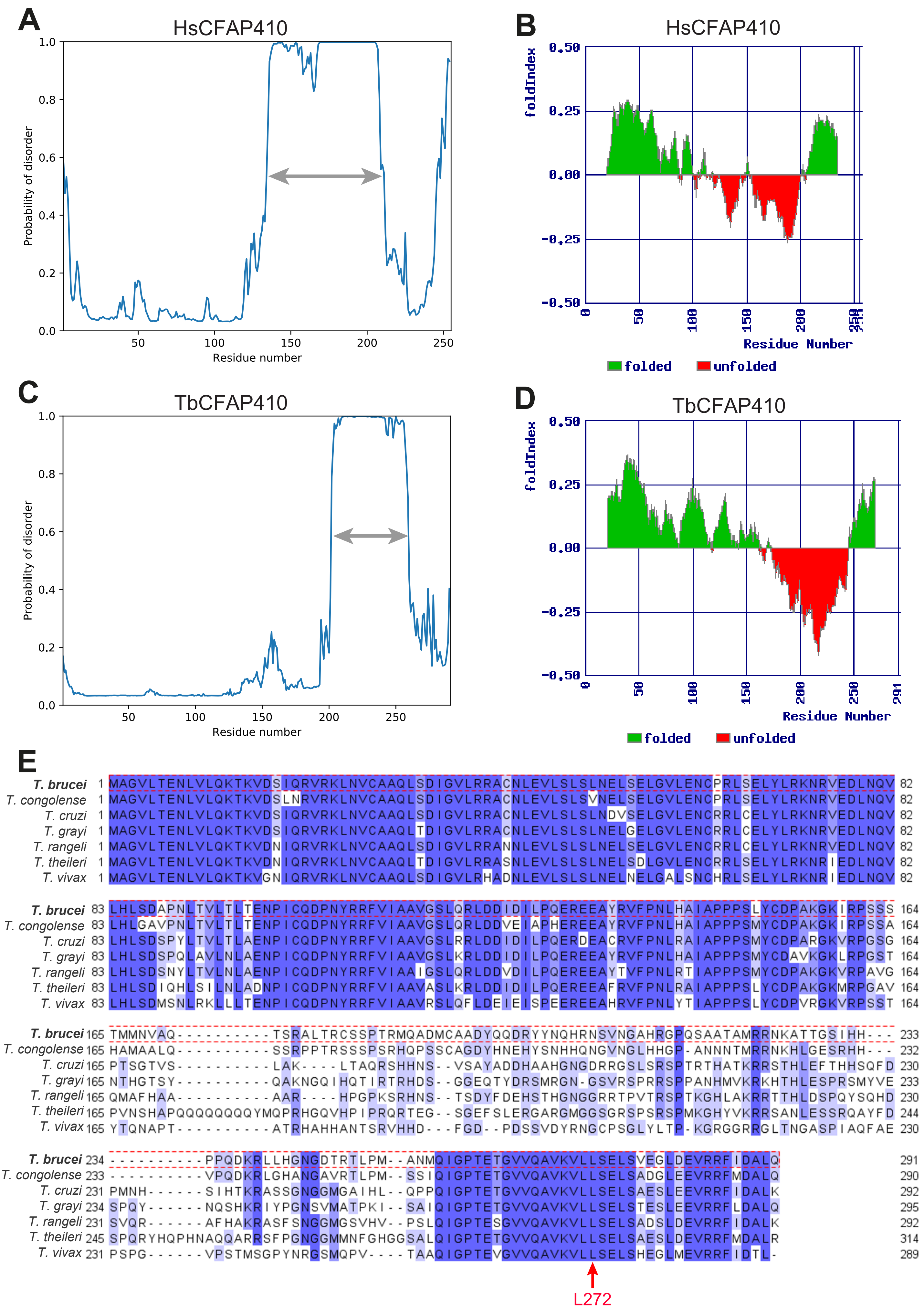
A
HsCFAP410
Probability of disorder
Residue number
B
HsCFAP410
foldIndex
Residue Number
folded
unfolded
C
TbCFAP410
Probability of disorder
Residue number
D
TbCFAP410
foldIndex
Residue Number
folded
unfolded
E
T. brucei
T. congolense
T. cruzi
T. grayi
T. rangeli
T. theileri
T. vivax
L272

Figure S2

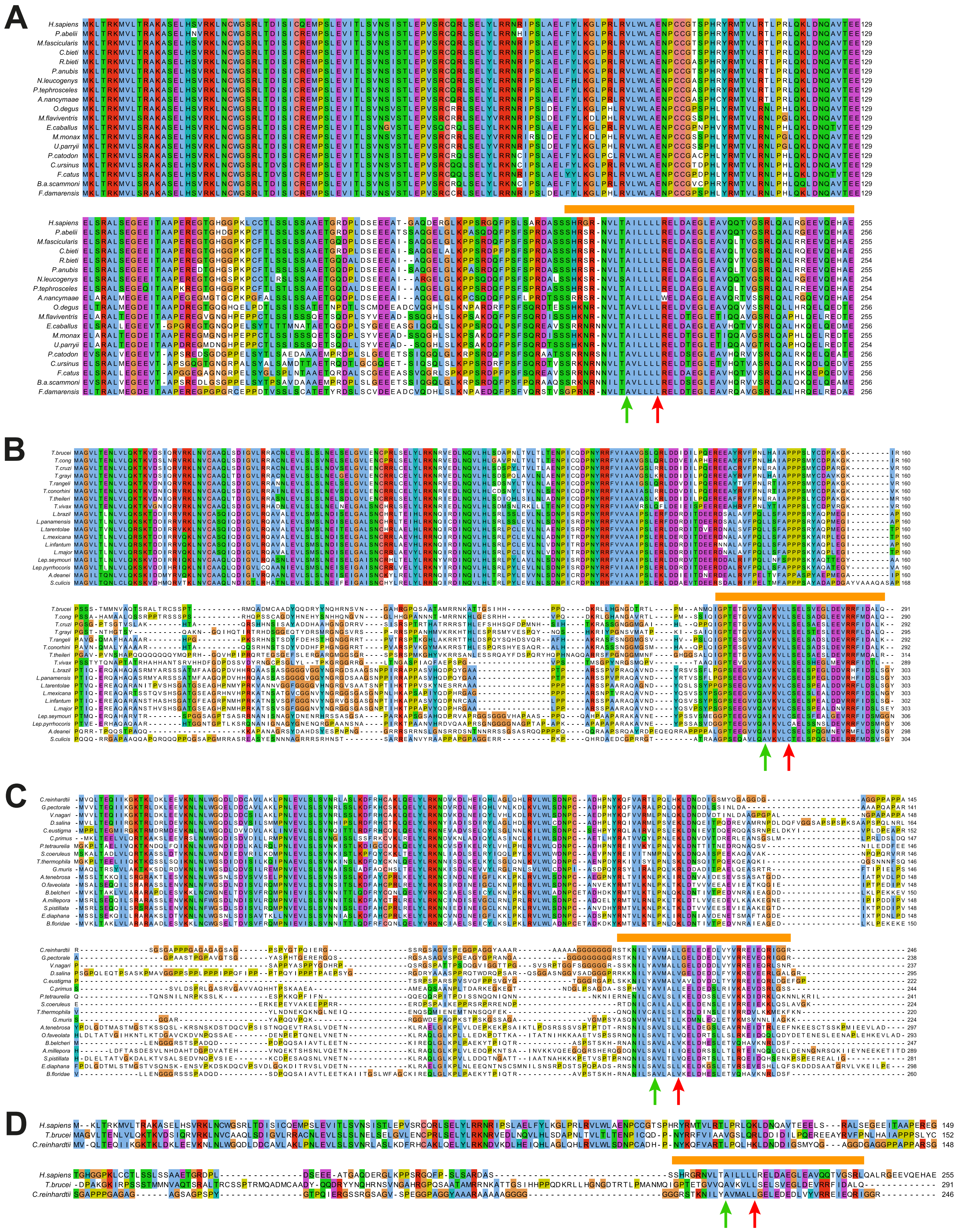
A
H.sapiens
P.abelii
M.fascicularis
C.bieti
R.bieti
P.anubis
N.leucogenys
P.tephrosceles
A.nancymaae
O.degus
M.flaviventris
E.caballus
M.monax
U.parryii
P.catodon
C.ursinus
F.catus
B.a.scammoni
F.damarensis
B
C
D
H.sapiens
T.brucei
C.reinhardtii

Figure S3

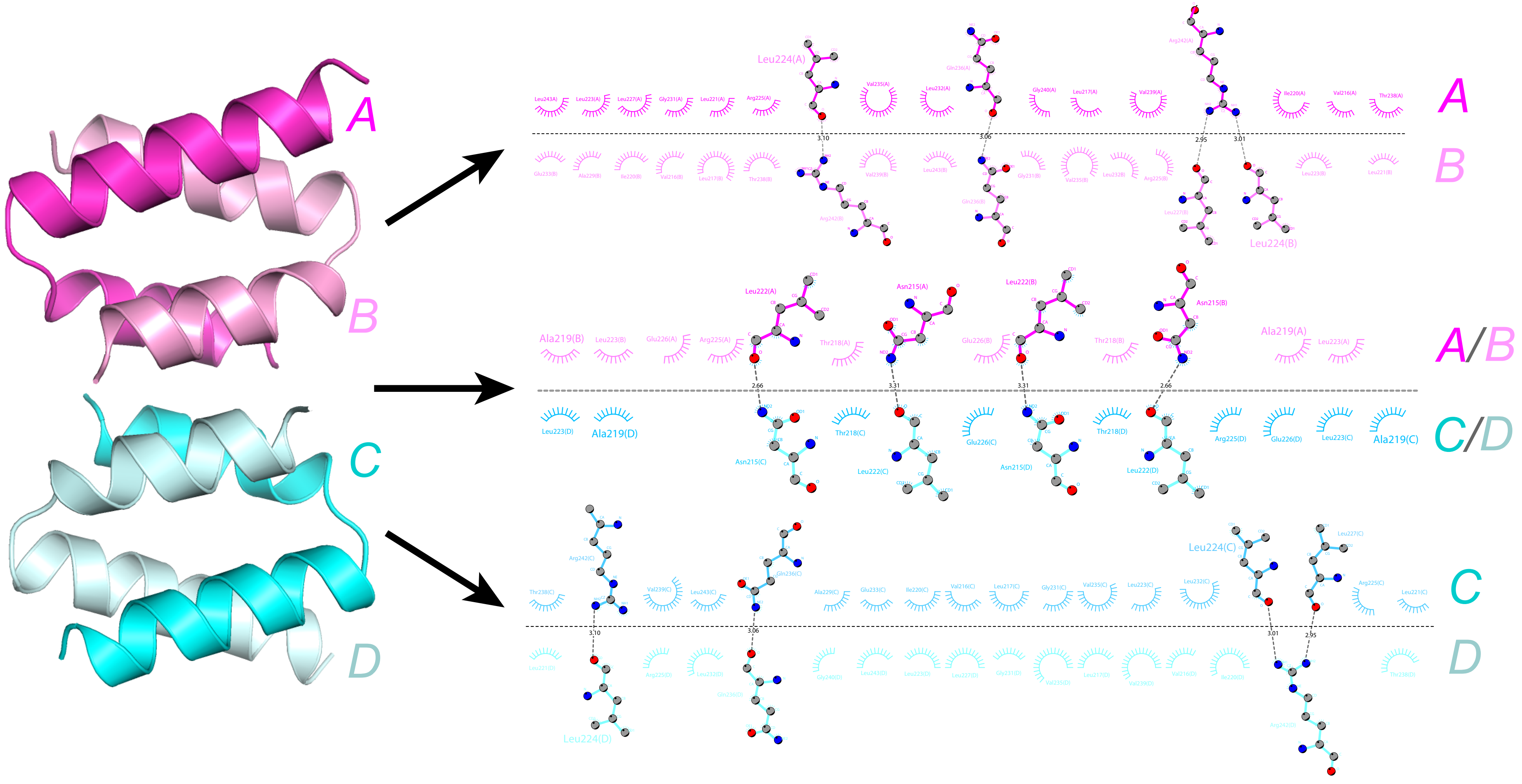
A
B
C
D
Leu224(A)
Leu224(B)
A
B
Leu222(A)
Asn215(A)
Leu222(B)
Asn215(B)
Ala219(B)
Ala219(A)
2.66
3.31
3.31
2.66
Ala219(D)
Ala219(C)
A/B
C/D
Leu224(C)
Leu224(D)
C
D
